# Teaching methods shape neural tuning to visual words in beginning readers

**DOI:** 10.1101/446203

**Authors:** Alice van de Walle de Ghelcke, Bruno Rossion, Christine Schiltz, Aliette Lochy

**Author notes:** Corresponding author: Aliette Lochy, UNIVERSITY OF LUXEMBOURG, Faculty of Language and Literature, Humanities, Arts and Education Research Unit Education, Culture, Cognition and Society Institute of Cognitive Science and Assessment, CAMPUS BELVAL, Maison des Sciences Humaines 11, Porte des Sciences L-4366 Esch-sur-Alzette Luxembourg.

## Abstract

The impact of global vs. phonics teaching methods for reading on the emergence of left hemisphere neural specialization for word recognition is unknown in children. We tested 42 first graders behaviorally and with electroencephalography with Fast Periodic Visual Stimulation to measure selective neural responses to letter strings. Letter strings were inserted periodically (1/5) in pseudofonts in 40sec sequences displayed at 6Hz and were either words globally taught at school, eliciting visual whole-word form recognition (global method), or control words/pseudowords eliciting grapheme-phoneme mappings (phonic method). Selective responses (F/5, 1.2Hz) were left lateralized for control stimuli but bilateral for globally taught words, especially in poor readers. These results show that global method instruction induces activation in the right hemisphere, involved in holistic processing and visual object recognition, rather than in the specialized left hemisphere for reading. Poor readers, given their difficulties in automatizing grapheme-phoneme mappings, mostly rely on this alternative inadequate strategy.

## Introduction

Reading is an essential prerequisite for the acquisition of knowledge across all school disciplines. It is also a complex skill, acquired only with formal instruction. Yet, little is known about how different teaching methods influence the development of neural circuits for reading. This lack of knowledge is surprising given its critical relevance for pedagogical and clinical purposes. The current practice in many schools is that first grade teachers use a so-called “mixed” approach, using two different methods in parallel for teaching to read. The “global” method requires children to visually memorize words globally and the phonic method teaches them letters-speech sounds mappings (grapheme-phoneme mappings, GP hereafter). In a natural school context, the present study assessed the cortical impact of these two different teaching methods in first grade children, by comparing neural responses to words that have been learnt globally to control letter strings (words/pseudowords) that constrain the use of GP mappings. In order to disentangle the potential role of familiarity by itself (intrinsically linked with the global method which involves an item-by-item learning), we then examined if responses to global words vary according to the mastery of GP mappings in two groups of children similarly exposed to global words.

Adults’ expert reading is characterized by highly automated recognition of written words. This automaticity allows accurate, effortless and fast (200 ms per word; Rayner, 1998 (Rayner, Pollatsek, & Schotter, 2012)) access to words representation integrating their orthographic, phonological and semantic properties. Neuroimaging studies have shown that this expertise relies on a left cortical network involving three circuits (ventral, dorsal, anterior) which are activated to interconnect vision and language. It is now admitted that a subregion of the left ventral occipito-temporal cortex (L-VOTC) termed the Visual Word Form Area (VWFA; Cohen et al., 2000) is crucial for the fast recognition of written words (see also (Lochy et al., 2018) for electrophysiological intracerebral evidence). Through its connections with phonological and lexico-semantic systems, VWFA triggers access to word’s phonological and semantic properties (Jobard, Crivello, & Tzourio-Mazoyer, 2003; McCandliss, Cohen, & Dehaene, 2003).

Before becoming fluent readers, children undergo laborious formal instruction during the first two years of primary school. Developmental studies revealed that the left circuits’ specialization for reading is driven by children’s early reading experience (Brem et al., 2010; Lochy, Van Reybroeck, & Rossion, 2016) and reading acquisition (Church, Coalson, Lugar, Petersen, & Schlaggar, 2008; Dehaene-Lambertz, Monzalvo, & Dehaene, 2018; Dundas, Plaut, & Behrmann, 2013; Eberhard-Moscicka, Jost, Raith, & Maurer, 2015; Gaillard, Balsamo, Ibrahim, Sachs, & Xu, 2003; Maurer et al., 2006; Pugh et al., 2001; Schlaggar & McCandliss, 2007; Turkeltaub, Gareau, Flowers, Zeffiro, & Eden, 2003). In line with the Phonological Mapping Hypothesis (Maurer & McCandliss, 2007), this left specialization is thought to emerge during the learning of GP mappings which induces progressive connections between posterior visual regions (letters representations) and anterior language regions (speech sounds representations).

The hypothesis that expert reading is built from the automatization of analytical processes performed on written words (e.g. orthographic, visuo-attentional, phonological) has been supported by developmental models (e.g. Ans, Carbonnel, & Valdois, 1998; Ehri, 1992; Frith, Patterson, Marshall, & Coltheart, 1985; Grainger, Lété, Bertand, Dufau, & Ziegler, 2012; Perfetti, 1991; Seymour, 1994). The acquisition of stable GP mappings and accurate knowledge of letters’ position in the word would be necessary conditions for the strengthening of words’ orthographic representation allowing its automated recognition (Perfetti, 1992). Indeed, several correct phonological recoding of a written word are necessary in order to store the word’s representation in the orthographic lexicon (self-teaching hypothesis; Cunningham, Perry, Stanovich, & Share, 2002; Share, 1995, 1999; Bowey & Muller, 2005; Nation, Angell, & Castles, 2007). Finally, training with GP mappings has been proposed to induce a refinement of phonological awareness which is known as a crucial predictor of reading acquisition (Byrne & Fielding-Barnsley, 1991; Goswami, 1993; Morais, Cary, Alegria, & Bertelson, 1979; Perfetti, Beck, Bell, & Hughes, 1987; Stuart & Coltheart, 1988; Vellutino & Scanlon, 1991).

At the earliest stages of teaching to read, many teachers use two different methods in parallel (“mixed” approach). The first method, which we refer to as “phonic”, involves explicit and progressive teaching of GP mappings, through a variety of exercises and items, allowing transfer on new letter strings. The second method, which we refer to as “global”, involves teaching of a strict visual recognition strategy (memorization of the visual shape of the whole word) aiming at creating a direct mapping between the written word, its spoken form, and its meaning. This method is characterized by a high number of repetitions of a few items inducing rote-learning. Initially, the global method (also termed “ideo-visual method”) claimed to develop expert skills (fast and accurate lexical recognition) independently of analytical skills. The aim was to allow children an immediate access to meaning, and therefore increase the pleasure of reading, without the slow and effortful decoding involved in GP mappings (Foucambert, 1976). Intense pedagogic debates persisted for years between approaches which emphasize learning the alphabetic code (e.g. phonic, syllabic method) and those emphasizing fast access to comprehension (e.g. global, ideo-visual, natural method) (Chall, 1967). Currently, the global method is no longer used in isolation, but rather in mixed approaches (Deauvieau & Terrail, 2018). According to surveys of Belgian and French 1st grade teachers, mixed approaches are the current dominant practice (between 60 and 90%) while strict alphabetical approaches are used by about only 10% of teachers (Nyssen & Lafontaine, 2006; Association Lire-Écrire, 2010 in Deauvieau, Espinoza, & Bruno, 2013). Critically, the potentially different neural mechanisms recruited by these teaching methods, have never been studied in children. To our knowledge, the impact of teaching methods in children has only been assessed behaviorally, while neural data are only available in adults and artificial learning contexts.

Behavioral classroom studies have shown that the type of method has a higher impact on children’s future reading performances than other variables (e.g. socio-economic background, performances in kindergarten, teachers’ experience) (Braibant & Gerard, 1996; Deauvieau et al., 2013; Goigoux, 2000). These studies and meta-analyses (Ehri, Nunes, Stahl, & Willows, 2001; Rayner, Foorman, Perfetti, Pesetsky, & Seidenberg, 2001) conclude that strictly alphabetical approaches are more effective than mixed or non-alphabetical approaches in word reading, spelling, and in text comprehension (Deauvieau & Terrail, 2018). Alphabetical approaches also induce the highest improvements in children at risk of a reading disorder or with low socio-economic background, by increasing their self-teaching ability (Ehri et al., 2001; Goigoux, 2016; Rayner et al., 2001). On the contrary, mixed or non-alphabetical approaches give rise to the highest proportion of poor readers and generate a higher heterogeneity of performance within a class (Braibant & Gerard, 1996; Deauvieau et al., 2013; Goigoux, 2000). In agreement with the self-teaching hypothesis (Share, 1995), sole visual exposure or limited phonological recoding (e.g. concurrent articulation) significantly reduces orthographic learning of novel letter strings (De Jong, Bitter, Van Setten, & Marinus, 2009; Kyte & Johnson, 2006; Share, 1999). Also, kindergarten children trained to memorize an artificial script or one-syllable words with a global strategy, have a lower ability to read novel stimuli than children trained with a GP mapping strategy (Jeffrey & Samuels, 1967) and have difficulties to infer GP mappings that have not been trained explicitly (Byrne, 1991, 1996).

Adults’ behavioral studies confirmed the lower efficiency of the global method concerning the transfer to novel stimuli and the implicit acquisition of GP mappings (Bishop, 1964; Bitan & Karni, 2003; Byrne, 1984; McCandliss, Schneider, & Smith, 1997; Yoncheva, Blau, Maurer, & McCandliss, 2010). In neuroimaging, fMRI training studies used unfamiliar stimuli (e.g. artificial script, pseudowords) to investigate neural correlates of orthographic, phonological and semantic processes in learning to read novel words. In comparison to global training (e.g. visual shape recognition, direct mapping with meaning), training based on phonological recoding led to a response modulation in the L-VOTC (Xue, Chen, Jin, & Dong, 2006; Sandak et al., 2004). Furthermore, the specific involvement of the VWFA in mapping letter shapes to phonology has been demonstrated in a study which contrasted the association of an artificial script with speech sounds and non-speech sounds (Hashimoto & Sakai, 2004). EEG studies showed that training GP mappings led to a left-lateralized N170 response sensitive to trained and untrained artificial script (between subjects design: Yoncheva et al., 2010); within-subjects design: Yoncheva et al., 2015), while global training led to a right-lateralized N170 response. These results highlight that the unit’s size on which the learner focuses, and therefore the processes engaged in word recognition after learning, directly influence brain mechanisms. It also suggests that only a phonic method engages the typical left lateralized brain circuitry of reading.

However, we currently do not know how (different) teaching methods impact brain activity in children. The current study aimed at filling this gap, by studying neural changes induced by the global and phonic teaching methods in a natural school context. We tested forty-four children (of whom 2 were excluded, see Material and Methods) during the first trimester of Grade 1. At that time of the school year, some words have been taught with a global method, in parallel to the teaching of GP mappings with a phonic method. This gave us the opportunity to compare neural responses to letter strings that should trigger only GP mappings (control words/pseudowords), to letter strings that should trigger visual whole form recognition processes (global words). The comparison of words globally learnt to control letter strings includes an inevitable and intrinsically confounded variable, which is familiarity (only global words are familiar to children). However, as explained in our hypotheses below, a differential impact of reading level on the use of a strict visual recognition strategy is expected, helping us to disentangle these confounded factors.

More precisely, in the two schools where we conducted the study, a list of global words was provided by the teachers, from which we extracted a common sub-part of twenty 4- and 5-letters words. These words have been globally taught in the classroom (mapping of whole-visual word form to spoken form, without knowledge of the inner letter-speech sounds associations), then printed on small cards given to the children in the so-called “words’ box”, and home-trained every day. In parallel, teachers taught GP mappings at the rate of one letter per week, with a variety of exercises: write the letter, recognize it within words, learn its case variants, etc. At the time of our testing, 9 letters had been taught in classrooms (a, é, e, è, I, o u, r, l). The other letters have been encountered visually by children and were variably recognized (as attested by performance in behavioral tests, see Table 1).

**Table 1.** Descriptive statistics for behavioral testing (N = 42)

| Behavioral tests and sub-tests | Scores |  |  |
| --- | --- | --- | --- |
|  | Min | Max | Mean (SD) |
| General cognitive functions |  |  |  |
| Nonverbal intelligence (CPM, accuracy in %) | 55.56 | 97.22 | 76.26 (12.20) |
| Selective attention (TEA-Ch, speed in sec) | 2.35 | 13.71 | 6.75 (2.36) |
| Vocabulary production (N-EEL, accuracy in %) | 56.14 | 94.74 | 77.63 (9.57) |
| Reading ability |  |  |  |
| Single letters (BELO, accuracy in %) | 19.23 | 100 | 71.43 (20.48) |
| Composite score (BELO, BALE, accuracy in %) | 2.40 | 74.87 | 34.57 (18.75) |

**Table 1-Source Data 1:** individual behavioral data summarized in Table 1 are provided as an excel file which is available through the following source data.

**Source data 1.** EEG data in frequency domain and behavioral data. https://doi.org/10.5061/dryad.0k6t3s0

We used a recently developed Fast Periodic Visual Stimulation (FPVS) approach with EEG recordings. In this approach, target stimuli are inserted periodically every five items within a rapid stream of base stimuli (6Hz) (Fig. 1), that were constituted of pseudofonts. Target stimuli (1 every 5, at 1.2Hz) were either words learned globally at school (GW) or control letter strings (words/pseudowords, W/PW). Given its high sensitivity (high Signal-to-Noise Ratio, SNR) (Norcia, Appelbaum, Ales, Cottereau, & Rossion, 2015; Regan, 1989), FPVS allows to rapidly (i.e. in a few minutes) measure selective responses to target stimuli. Recently, robust discrimination response for words among pseudofonts, non-words, or even pseudowords was shown in adults over the left occipito-temporal cortex (Lochy, Van Belle, & Rossion, 2015). In 5 years-old preschoolers, a discrimination response for letter strings (words/pseudowords) within pseudofonts was observed over left posterior regions of the scalp and, critically, correlated with GP knowledge (Lochy et al., 2016). These findings revealed the potential of the FPVS-EEG approach to assess the neuro-cognitive representations of written words and the development of neural circuits for reading (see also Lochy et al., 2018 for intracerebral evidence). Furthermore, as a behavior-free (implicit and automatic visual discrimination of letter strings) and rapid approach, FPVS-EEG is ideal to use in young children.

**Fig. 1.**
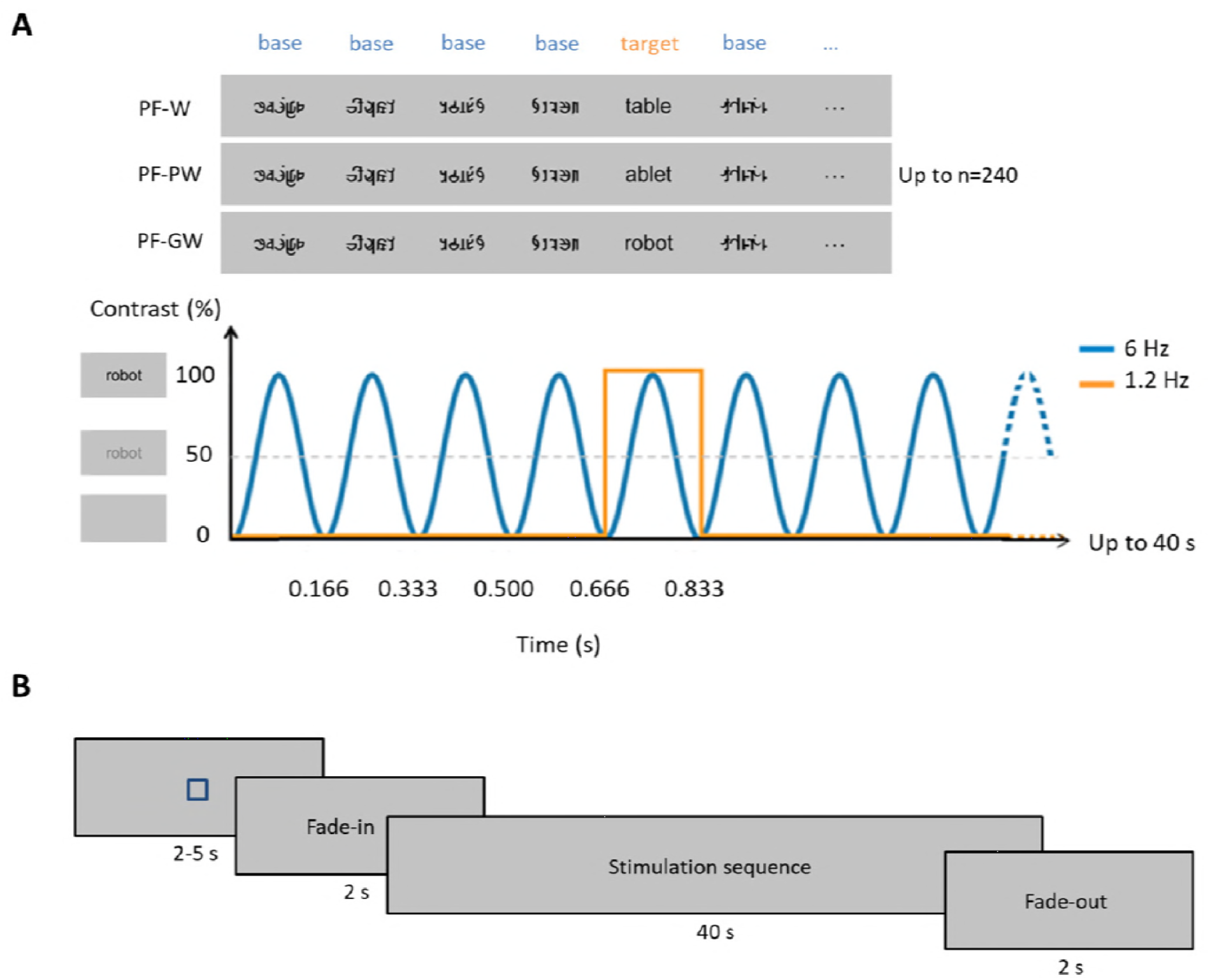
Experimental paradigm. **(A)** In each condition, base stimuli were pseudofonts (top, middle and bottom rows), and target stimuli were either control words (W, top row), pseudowords (PW, middle row) or globally taught words (GW, bottom row) appearing every fifth item. Each sequence lasted 40 seconds, during which stimuli were presented by sinusoidal contrast modulation at 6 Hz, each stimulus reaching full contrast after 83 ms (i.e., one cycle duration = 166.66 ms). Stimulation alternated between base (B) and target (T) stimuli such as BBBBTBBBBTBBB. Target stimuli therefore appeared at 6Hz/5, so at 1.2Hz. Stimuli were randomly presented with no immediate repetition and appeared continuously on the screen. In total, 240 stimuli were presented per sequence (48 target stimuli and 192 base stimuli), and each condition was repeated 3 times. **(B)** Timeline of a sequence: each sequence started with a fixation square (for 2-5 s) after which the stimulation faded in (for 2 s) then reached full contrast (for 40 s) and then faded out (for 2 s); see Methods.

In the current study, we tested the hypothesis that, at an early stage of reading acquisition, words learnt globally as unique visual forms would be represented differently than control letter strings (words/pseudowords) which would trigger GP mappings. More specifically and based on our previous work in kindergarten children (Lochy et al., 2016), we expected that responses at 1.2Hz and harmonics, reflecting discrimination of letters among pseudofonts, would vary according to the condition. We hypothesized that GP mappings (in W/PW control letter strings) would engage the typical left brain circuitry of reading, while GW would engage a right-lateralized circuitry (Yoncheva et al., 2010, 2015). Most importantly, we expected a differential effect of the global teaching method as a function of children’s reading level, which if confirmed also precludes interpretation of our results as simple familiarity effect. Indeed, poor readers having difficulties to automatize GP mappings (e.g. unstable knowledge of letters’ sounds (Perfetti, 1992), deficit in phonological awareness (Ziegler et al., 2010) and/or visuo-attentional processing (Ans et al., 1998)), compensate their difficulties by using alternative strategies for reading, like visual memory among others (e.g. salient visual features; Campbell & Butterworth, 1985; Vellutino, 1987). Therefore, we hypothesized that they might be more, or exclusively, influenced by the global method inciting them to visually encode the whole-word forms. In other terms, they might show a different response pattern to GW than good readers (e.g. more right-lateralized). Because children of both groups have been familiarized explicitly and to the same degree with the GW, group differences regarding responses to GW would strongly suggest that familiarity per se is not the key factor inducing a right-lateralized neural response. Indeed, if familiarity induces engagement of the right hemisphere, then this should be the case whatever the children’s reading level.

## Results

### 1. Whole group analysis

#### 1.1. Word discrimination responses

Selective discrimination responses to letter-strings (W, PW, GW) inserted in pseudofonts were significant (Z-score > 2.58) at 1.2Hz and several harmonics (F/5 to 4F/5) at nine posterior electrodes (O1, O2, Oz, P7, P8, PO3, PO4, P3, P4). Electrodes were ranked according to their largest amplitude values for the sum of baseline corrected amplitudes computed on four significant harmonics (1.2Hz, 2.4Hz, 3.6Hz, 4.8Hz, see Methods) as determined by grand-averaged data. In all conditions, most of the response was captured on the left occipital channel O1 (= 2.35 μV), the signal dropping rapidly on nearby electrodes (P7 = 1.73 μV; PO3 = 1.07 μV; P3 = 0.13 μV). Therefore, in line with our previous study in children (Lochy et al., 2016), we focused on O1 and its homologous right hemispheric electrode O2.

An ANOVA was performed on target discrimination responses (sum of baseline corrected amplitudes) with *Hemisphere* (left-O1, right-O2) and *Condition* (PF-W, PF-PW, PF-GW) as within-subjects factors. It revealed a main effect of *Hemisphere* [F_1,41_ = 9.24, *P* = 0.004, η^2^ = .20], no effect of *Condition* [F_2,82_ =0.82, *P* = 0.445, η^2^ = .02] and a trend for a significant interaction between these two factors [F2,82 = 3.08, *P* = 0.051, η2 = .07]. Paired samples t-tests were performed in order to compare the response amplitude between the left (O1) and the right (O2) hemispheres in each condition. Response amplitude to both control words (O1 = 2.31 μV, O2 = 1.44 μV) and pseudowords (O1 = 2.37 μV, O2 = 1.64 μV) was stronger in the left (O1) than in the right (O2) (PF-W: [*t*(41) = 3.45; *P* = 0.001]; PF-PW: [*t*(41) = 3.16; P = 0.003]), but response amplitude to global words did not significantly differ between O1 (2.37 μV) and O2 (1.97 μV) (PF-GW: ([*t*(41) = 1.59; *P* = 0.120]) (Fig.2).

#### 1.2. Base rate responses

The base stimulation frequency reflects the general synchronization of the visual system to the periodic stimulation (or level of attention to the stimuli). Z scores computation revealed significant responses in all conditions at exactly 6Hz and several harmonics (F to 5F) at six medial occipito-parietal electrodes (O1, O2, Oz, PO3, PO4, Pz). In order to determine the electrodes of interest, all electrodes were ranked according to their largest amplitude values for the sum of baseline corrected amplitudes computed on five significant harmonics (6Hz, 12Hz, 18Hz, 24Hz, 30Hz, see Methods) as determined by grand averaged data. In all conditions, the largest response was recorded at three occipito-medial (OM) electrodes: O1 (2.72 μV), O2 (3.04 μV) and Oz (3.18 μV). Therefore, we averaged their amplitude values for analyses (OM ROI = mean O1, O2, Oz) (Fig.2). An ANOVA performed on response amplitudes in OM ROI with *Condition* (PF-W, PF-PW, PF-GW) as within-subjects factor, did not reveal any effect of *Condition* [F_2,82_ = 1.62, *P =* 0.205, η^2^ = .04].

**Fig. 2.**
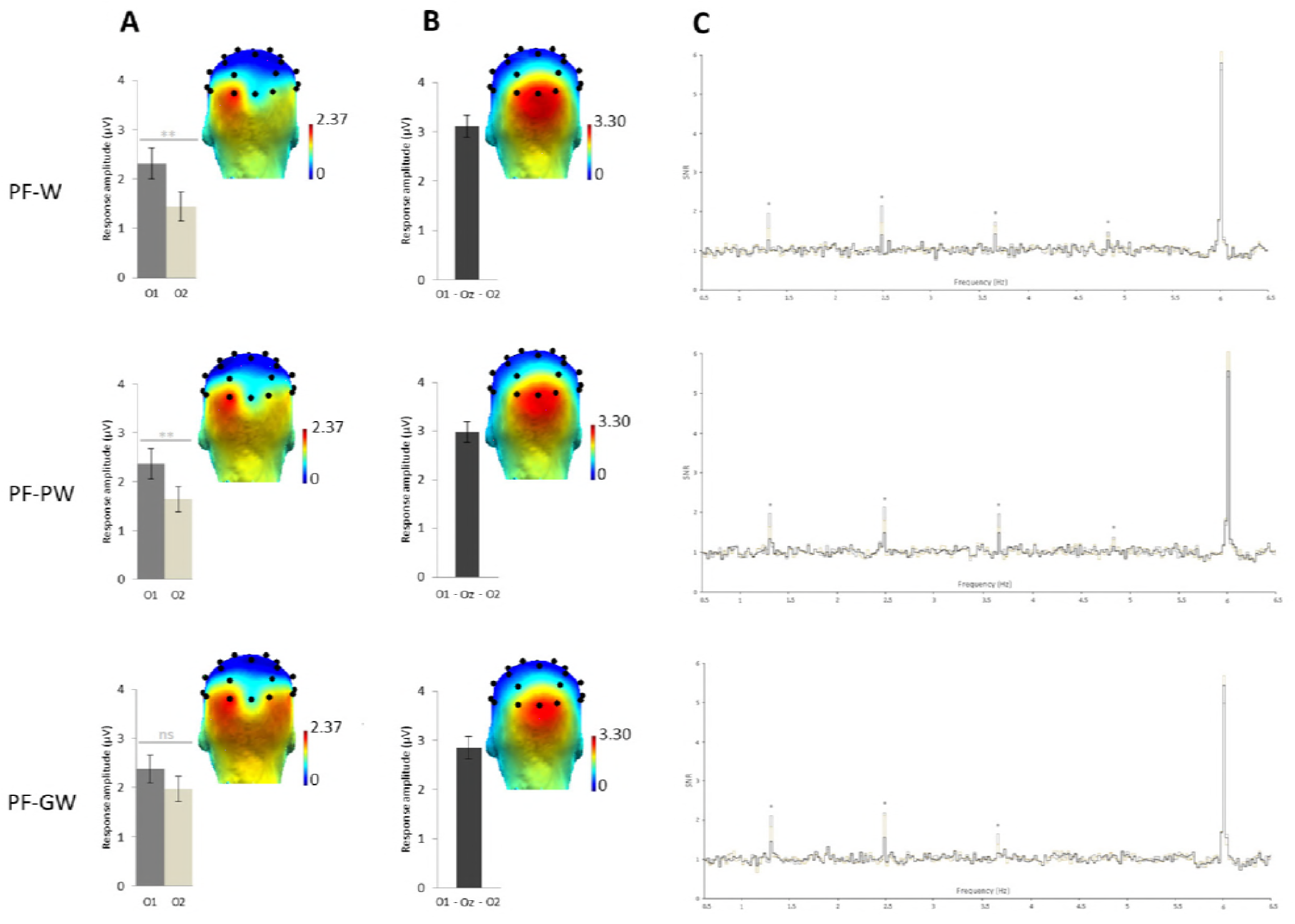
Response amplitudes and topographies of the whole group. Grand-averaged (N=42) scalp topographies for the response in each condition at **(A)** the target and **(B)** base frequencies (sum of baseline subtracted amplitudes at significant harmonics; see Methods). Histograms represent the same data, with standard errors of the mean. For letter strings-selective responses, stars indicate significant difference between the left (O1) and the right (O2) occipital electrodes (**: p<0.01, ns: not significant). **(C)** SNR EEG spectra on O1 (dark grey), O2 (in light grey) and Oz (in black) for each condition, stars indicate significant responses at letter strings-selective frequency and harmonics (1.2Hz, 2.4Hz, 3.6Hz, 4.8Hz). **Figure 2-Source Data 1**. Individual EEG data summarized in Figure 2 are provided as a compressed folder which is available through the following source data. **Source data 1.** EEG data in frequency domain and behavioral data. https://doi.org/10.5061/drvad.0k6t3s0

### 2. Analysis by reading level

First, we computed a composite score of reading for each child by averaging accuracy scores for graphemes, syllables and words reading. Since our objective was to compare subgroups, we assigned children in subgroups on the basis of the group’s mean composite score (34.57% of accuracy). Children who performed above the group’s mean composite score (> 35%) were considered as « good readers » (N=18), and those below (< 34%) as « poor readers » (N=19). We excluded from analysis 5 children whose score was at the group’s mean composite score (34-35%).

#### 2.1. Word-discrimination responses

An ANOVA was performed on word discrimination responses (sum of baseline corrected amplitudes) with *Hemisphere* (left-O1, right-O2) and *Condition* (PF-W, PF-PW, PF-GW) as within-subjects factors and *Group* (good readers, poor readers) as between-subjects factor. It revealed an effect of *Hemisphere* [F_1,35_ = 18.38, *P* = 0.000, η^2^ = .34], responses being overall stronger in the LH (2.41 μV vs. 1.59 μV in the LH and RH respectively), an interaction between *Hemisphere* and *Group* [F_1,35_ = 4.96, *P* = 0.033, η^2^ = .12], responses of the two groups were very similar in the RH (good readers: 1.60 μV, poor readers: 1.57μV), while good readers had a stronger response than poor readers in the LH (2.84 μV and 2.01 μV respectively) and a trend for interaction between *Hemisphere* and *Condition* [F_2,70_ = 2.89, *P* = 0.063, η^2^ = .08] but no main effect of *Condition, Group* or any other interaction (all Fs < 1). Given the interactions between *Hemisphere* × *Group* and *Hemisphere* × *Condition*, and the trends highlighted by the topographies and the histograms (Fig. 3), paired t-tests were performed between O1 and O2 in each condition and group. Good readers presented a significantly stronger response in the left hemisphere in each condition; PF-W: [*t*(17) = 4.53; *P* = 0.000] (O1 = 2.86 μV, O2 = 1.29 μV), PF-PW: [*t*(17) = 3.67; *P* = 0.002] (O1 = 2.84 μV, O2 = 1.66 μV), PF-GW: [*t*(17) = 2.93; *P* = 0.009] (O1 = 2.82 μV, O2 = 1.78 μV). Poor readers presented a trend for left lateralized response in PF-W [*t*(18) = 1.80; *P* = 0.088] (O1 = 1.98 μV, O2 = 1.42 μV), a left lateralized response in PF-PW [*t*(18) = 2.21; *P* = 0.040] (O1 = 1.92 μV, O2 = 1.38 μV) and a bilateral response in PF-GW [*t*(18) = 0.26; *P* = 0.802] (O1 = 2.12 μV, O2 = 2.03 μV).

#### 2.2. Base rate responses

An ANOVA performed on response amplitudes in OM ROI with *Condition* (PF-W, PF-PW, PF-GW) as within-subjects factor and *Group* (good readers, poor readers) as between-subject factor, revealed no effect of *Condition* [F_2,70_ = 1.51, *P* = 0.228, η2 = .04], no effect of *Group* [F_1,35_ = 3.39, *P* = 0.074, η^2^ = .09] and no interaction between these two factors [F_2,70_ = 2.46, *P* = 0.093, η^2^ = .07].

**Fig. 3.**
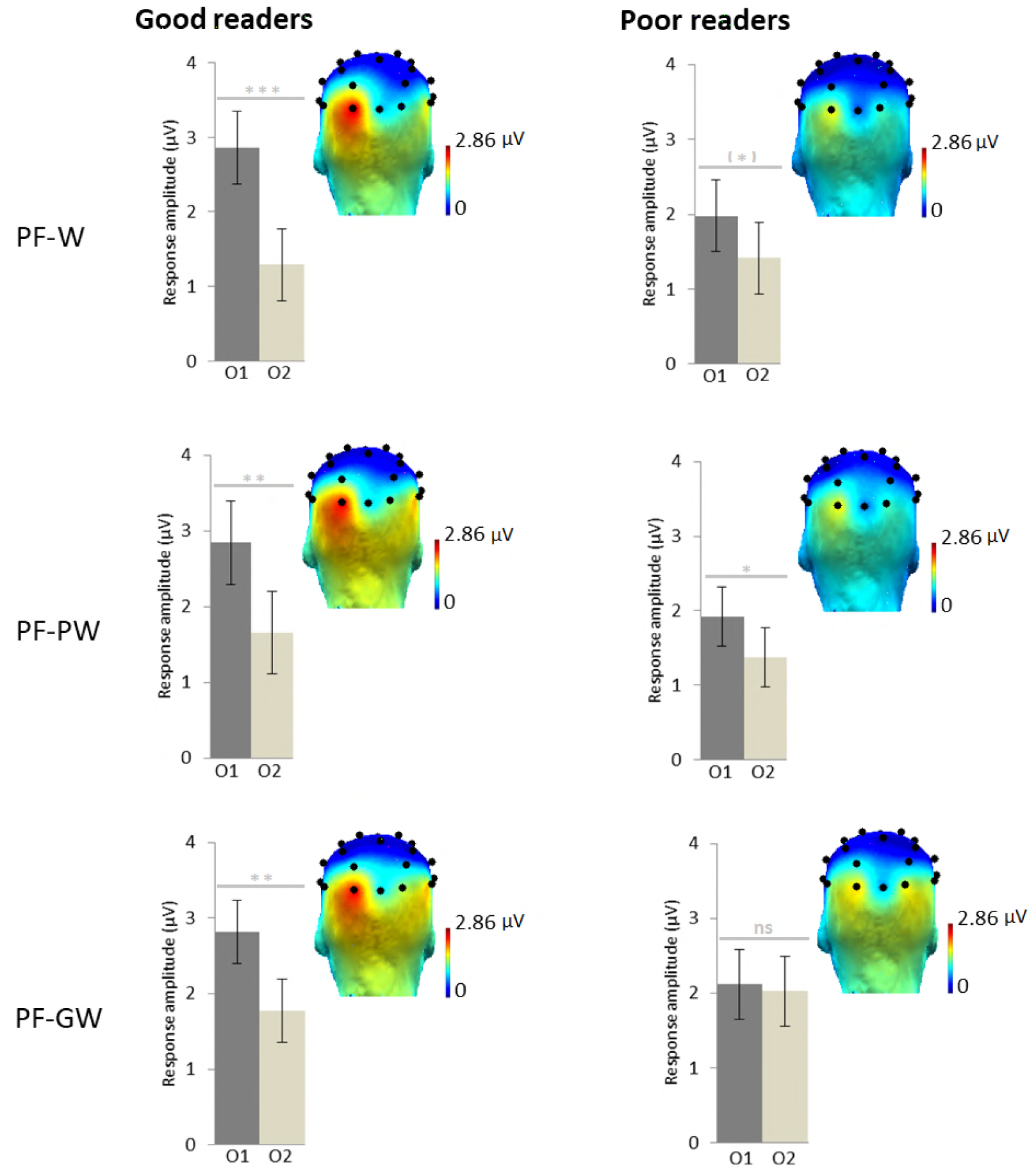
Response amplitudes and topographies by reading level. Scalp topographies for the response in each condition at the letter strings-discrimination frequency (sum of baseline subtracted amplitudes at significant harmonics: 1.2 Hz, 2.4 Hz, 3.6 Hz, 4.8 Hz; see Methods) for children assigned in the good readers group (N=18) and in the poor readers group (N=19) on the basis of their behavioral reading scores. Histograms represent the same data, with standard errors of the mean. Stars indicate significant difference between the left (O1) and the right (O2) occipital electrodes (*: p<0.05; **: p<0.01; ***: p<0.001, ns: not significant) **Figure 3-Source Data 1**. Individual EEG data summarized in Figure 3 are provided as a compressed folder which is available through the following source data. **Source data 1.** EEG data in frequency domain and behavioral data. https://doi.org/10.5061/dryad.0k6t3s0

#### 2.3. Brain-behavior correlations

We assessed if reading scores correlated with lateralization scores (LS; calculated as LH - RH) for the responses to letter strings on the 37 children retained in our subgroups analysis (Fig. 4), and this, separately for the response to global words, control words, control pseudowords, as well as the average of W/PW (as in Lochy et al., 2016). All these correlations were significant (respectively: Spearman Rho=0.44, *P* = 0.003; Rho=0.43, *P* < 0.004; Rho=0.29, *P* = 0.043; Rho=0.36, *P* = 0.016), indicating that better readers tended to reveal more left-lateralized responses in all conditions.

**Fig. 4.**
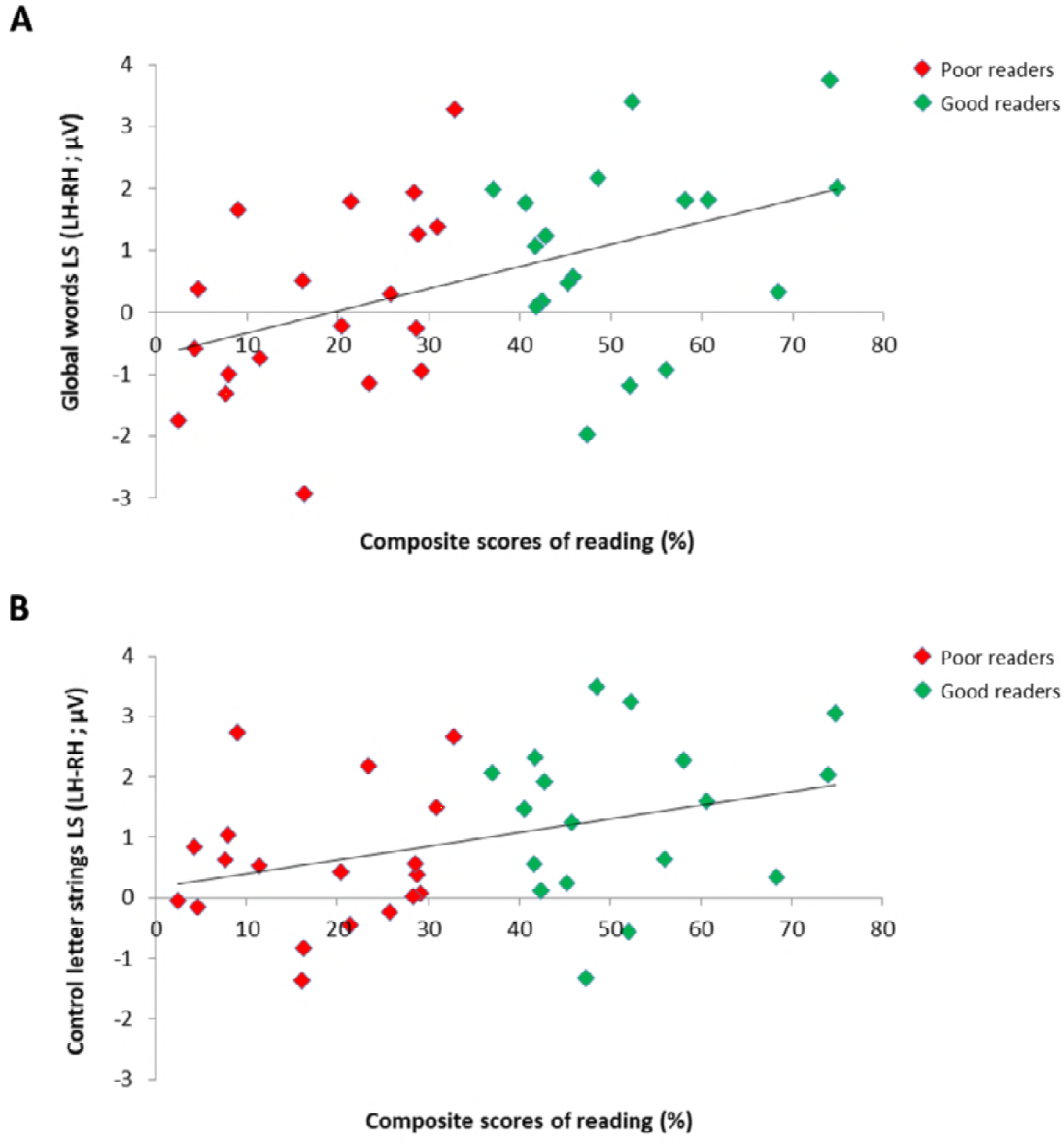
Relation between individual reading scores and EEG lateralization scores for the responses to letter strings in 37 children. Scatter plots of significant positive correlation between composite scores of reading (averaged accuracy scores for graphemes, syllables and words) and EEG lateralization scores (LH-RH) for the responses to **(A)** global words (Spearman Rho=0.44) and **(B)** control letter strings (averaged response to control words/pseudowords; Spearman Rho=0.36) in good readers (N = 18; green dots) and poor readers (N = 19; red dots).

Per group, we computed correlations between conditions, reasoning that if distinct processes are triggered for global words, then LS should not correlate with the other conditions. In the good readers group, all conditions correlated highly: W and PW (Spearman Rho=0.79; *P* = 0.000), W and GW (Spearman Rho=0.78; *P* = 0.000), PW and GW (Spearman Rho=0.76; *P* = 0.000). In the poor readers group, only W and PW correlated significantly (Spearman Rho=0.64; *P* = 0.002), while GW did not correlate with the two other conditions (with W and PW, respectively Spearman Rho=0.19; *P* = 0.22; and Rho=0.035; *P* = 0.44) (Fig. 5).

**Fig. 5.**
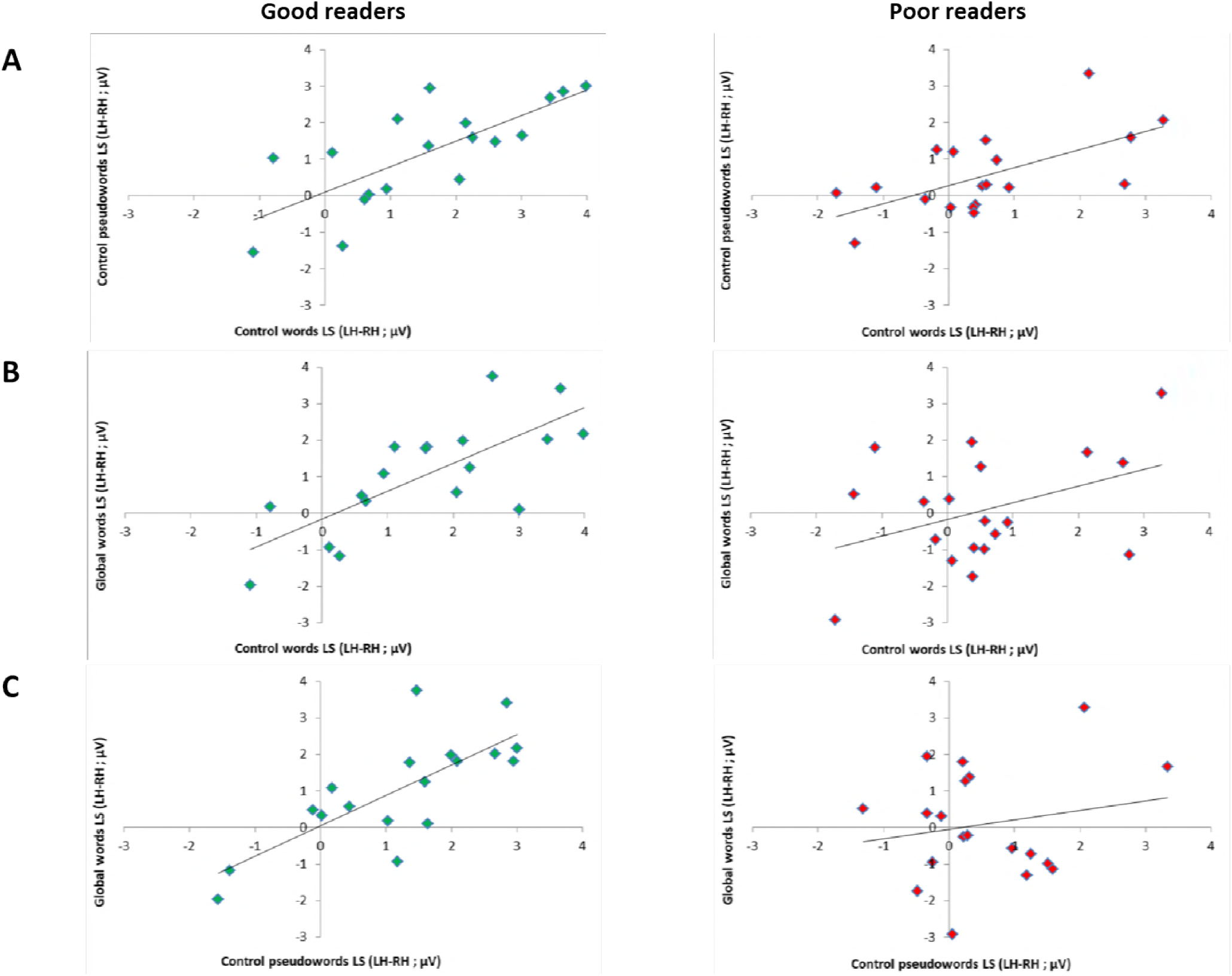
Relation between EEG lateralization scores (LS) across conditions per reading level subgroup. Scatter plots of correlation analysis between EEG lateralization scores (LH-RH) for the responses to **(A)** control words and pseudowords, **(B)** global words and control words and to **(C)** global words and pseudowords in good readers (N=18, green dots) and poor readers (N=19, red dots). Correlations were all highly significant in good readers (all p<0.001), while in poor readers, responses to global words did not correlate with control letter strings (with W: *P* = 0.22; with PW: *P* = 0.44), which correlated together (*P* = 0.002), suggesting atypical processing of global words.

## Discussion

Our study reveals that the methods used to teach children how to read impact the development of neural responses to letter strings. We took advantage of a natural school context using a mixed approach in the beginning of the first grade to compare, in 42 children, neural responses to letter strings that trigger GP mappings (phonic method), with letter strings that induce whole-word visual form recognition processes (global method) in a within-subjects design. In 2 minutes of recording only, we found different and specific neural responses to these two categories of letter strings, which were moreover modulated by the reading level of children. These results lead to two critical conclusions that will be discussed in turn.

First, GP mappings predominantly engaged the LH while whole-word recognition engaged also the RH. This confirms in children previous findings in adults trained with an artificial script, where globally learnt stimuli engaged the RH more than items learnt with a phonic approach (Yoncheva et al., 2010, 2015). At the behavioral level, developmental models support the hypothesis that expert reading skills are built from the automatization of analytical processes performed on written words (e.g. Ans et al., 1998; Ehri, 1992; Frith et al., 1985; Grainger et al., 2012; Perfetti, 1991; Seymour, 1994). Furthermore, in comparison to non-strict alphabetical instruction, strict alphabetical instruction has been identified as the most effective in both adults (Bishop, 1964; Bitan & Karni, 2003; Byrne, 1984; McCandliss et al., 1997; Yoncheva et al., 2010) and children (Braibant & Gerard, 1996; Byrne, 1991, 1996; De Jong et al., 2009; Deauvieau et al., 2013; Ehri et al., 2001; Goigoux, 2000; Goigoux, Cèbe, & Pironom, 2016; Jeffrey & Samuels, 1967; Kyte & Johnson, 2006; Rayner et al., 2001; Share, 1999). Here we show that the type of teaching method used, and hence the type of process triggered for reading, differentially influence the neural circuits activated when processing letter strings.

The left lateralization in response to letter strings is in line with previous developmental studies assessing brain responses to written material during reading acquisition (Brem et al., 2010; Church et al., 2008; Dehaene-Lambertz et al., 2018; Dundas et al., 2013; Eberhard-Moscicka et al., 2015; Gaillard et al., 2003; Maurer et al., 2006; Pugh et al., 2001; Schlaggar et al., 2002; Turkeltaub et al., 2003), including our previous study using the same FPVS-EEG approach in kindergarten children (Lochy et al., 2016). It supports the Phonological Mapping Hypothesis (Maurer & McCandliss, 2007) proposing that simple passive viewing of letter strings triggers activation of associations between orthographic and phonological representations. According to neuroimaging studies in adults, the dominant LH engagement is the signature of expert reading (Bentin, Mouchetant-Rostaing, Giard, Echallier, & Pernier, 1999; Cohen & Dehaene, 2004; Cohen et al., 2000, 2002; Urs Maurer, Zevin, & McCandliss, 2008; B. McCandliss et al., 2003; Petersen, Fox, Posner, Mintun, & Raichle, 1988), and responses increase in the L-VOT with age and reading level (Ben-Shachar, Dougherty, Deutsch, & Wandell, 2011; Brem et al., 2010; Maurer, Blau, Yoncheva, & McCandliss, 2010). Remarkably, the knowledge of all letters is not necessary to observe this LH lateralization pattern in response to letter strings. In the current study, control words/pseudowords were constituted of 18 different letters, of which only 9 had been formally taught with the phonic method at school, and on average 14 were known (see Table 1). Similarly, in our previous kindergarten study (Lochy et al., 2016), children who knew more than 9 letters already showed this typical LH pattern of responses to letter strings.

The RH, on the contrary, is known to support object and face recognition (Bentin, Allison, Puce, Perez, & McCarthy, 1996; Rossion, Joyce, Cottrell, & Tarr, 2003; Tanaka & Curran, 2001), and has been suggested to reflect visual familiarity with script in tasks where a minimal visual recognition strategy was possible, for instance in a one-back task (Maurer et al., 2010). At a broader level, the two hemispheres are hypothesized to preferentially support different types of processes, the LH showing an advantage for analytic/local processes while the RH shows an advantage for holistic/global processes (Corballis, 2003; Robertson & Ivry, 2000; Sergent, 1982), as evidenced by neuropsychological deficits of patients with unilateral brain damage (review in Ivry & Robertson, 1998). Our results reveal that globally learnt words enhance RH responses, which we interpret as a greater involvement of purely visual processes in their recognition. This atypical neural pattern during the processing of these letter strings suggests that whole-word instruction exposes children to a deviant (i.e., not optimal) reading strategy.

As children were tested after only 2-3 months of reading instruction, our results highlight a rapid impact of formal instruction on neural processes recruited during reading. Such a rapid influence of instruction has been previously reported in neuroimaging training studies in both children (e.g., after 8 weeks in Brem et al., 2010; 4 weeks in James, 2010) and adults (40 hours of training over 4 weeks in Xue et al., 2006); one session of 40 minutes in Yoncheva et al., 2010; 2 × 50 min over 2 days in Yoncheva et al., 2015).

As in our previous study in pre-readers (Lochy et al., 2016), and in recent fMRI findings (Dehaene-Lambertz et al., 2018), the left lateralized response to letter strings suggests an earlier emergence of the neural specialization for reading than previously found in EEG studies, where it was hypothesized to emerge after 1-1.5 year of formal instruction (Eberhard-Moscicka et al., 2015; Maurer et al., 2006). The correlation found here between reading scores and lateralization scores (Rho=0.36) is in the range of our previous data with kindergartners (Lochy et al., 2016; Lochy, de Heering, & Rossion, 2017, Rho=0.33 for letter production and 0.42 for letter recognition). Furthermore, as reading scores were assessed here with other behavioral tests than in our previous study (single letters reading in kindergarten vs. letters, syllables, words and pseudowords here), we can conclude that the relationship between response amplitude to letter strings measured in FPVS and performance at the beginning of the learning process, can be generalized to other age-groups and other behavioral tests.

Our second crucial finding is that poor readers were more influenced by the global teaching method than good readers. In fact, only the poor readers showed the atypical bilateral response to global words. This finding agrees with behavioral training studies showing that sufficient GP mappings knowledge is necessary for being able to infer GP mappings from globally trained words (Byrne, 1984, 1991, 1996) or even to switch from global to local attentional focus (Yoncheva et al., 2010, 2015). This means that good readers, having sufficiently automatized GP mappings, could process even the global words with an orthography-to-phonology type of process while poor readers relied on whole-word recognition. This finding supports the view that non-alphabetical approaches are significantly less efficient in children with learning difficulties (Braibant & Gerard, 1996; Deauvieau et al., 2013; Ehri et al., 2001; Goigoux, 2000; Rayner et al., 2001). Indeed, the global method may reinforce the use of non-efficient compensatory strategies already set up by poor readers (e.g. reading by guessing from semantic or visual clues). Furthermore, if GP mappings cannot be applied, the global method leads to a non-economical storage of written words comparable to visual objects and thus, may strongly interfere on reading accuracy (e.g. confusion between visually similar words) and reading acquisition in general, for instance by impeding self-teaching of novel words (Share, 1995), as well as development of phonological awareness and letter knowledge. The fact that good readers processed all stimuli types predominantly with their LH nevertheless shows the dominance of the most efficient process, once acquired.

The phonic and global methods that we compared differ by definition on an important aspect which is item familiarity. Indeed, the phonic method aims at providing the ability to transfer GP mappings on new words, while the global method aims at creating representations for the learnt items. Therefore, an intrinsic confounded factor inherent to the two methods is that global words were highly familiar to children, while control words and pseudowords have not been explicitly taught at school. However, the RH engagement for global words is unlikely to be due to familiarity as such. Indeed, all children were as familiar to the set of global words, but good readers presented a left lateralized response in all conditions, while poor readers presented RH engagement for global words only. These bygroup and by-condition differences confirm that global words are processed differently than control letter strings in poor readers only, and are not simply more familiar. The inter-conditions correlations by group provide another argument supporting the view that in good readers, the reading processes triggered by both control letter strings and global words were similar, while for poor readers, they were different. For good readers, the lateralization scores correlated highly between all three conditions (above Rho=0.75), while for poor readers, correlations between the two control letter strings (words/pseudowords) were significant (Rho=0.64), while they were not between global words and control letter strings. The RH engagement for global words is also unlikely to be due to a fluctuation of attention given that, first, base rate responses (i.e., 6Hz and harmonics) were similar across conditions and groups; and second, detection of the color change on the fixation square did not vary according to conditions and groups. All these findings thus confirm the specificity of the processes triggered by global words, which are different than the GP mappings triggered by control letter strings, and which are modulated by the mastery of GP mappings.

Since the persistence of the effects has not been assessed, this study does not allow us to conclude that the global method has a negative impact on the long-term development of neural circuits for reading. It may well be that before acquiring the mastery and automatization of GP mappings, good readers presented the same atypical neural pattern than poor readers for global words. Thus, it is also possible that poor readers, after improvement of GP mappings ability with formal instruction, could progressively present a left lateralized response despite the whole-words instruction. On the other hand, some children having a severe or specific disorder and unable to reach a sufficient level of GP mappings automatization, could maintain an atypical neural pattern for these global words. Furthermore, as suggested by behavioral studies (Campbell & Butterworth, 1985; Vellutino, 1987), they could even mostly attempt to transfer this strategy to process other written words, in order to compensate their persistent difficulties in acquiring GP mappings.

Therefore, the atypical neural circuitry that we find here for global words and only in poor readers, seem to us sufficient to strongly encourage education professionals not to use the global method for teaching to read, but on the contrary, to automatize GP mappings with alphabetical approaches. Visual memorization of global word shape is not involved in (adult) expert recognition, which results from the automatization of analytical processes performed on written words. We are aware that the aim of teachers is to vary the approaches in learning to read in order to motivate children, because the process of learning all the GP mappings and to automatize them is long and laborious. However, only strict alphabetical approaches can provide the indispensable foundations for the development of expert reading skills, while non-strict alphabetical approaches penalize pupils’ learning and this effect is probably even enhanced in children with learning difficulties.

## Conclusion

To our knowledge, our study is the first to assess the impact of two widely used teaching methods, global and phonic, on the neural circuits for reading in children. We showed, in a natural school context and at an early stage of reading acquisition, that the unit’s size involved in mapping written information to phonological forms (graphemes to phonemes vs visual whole-words to its spoken form) directly influences neural mechanisms recruited during written word processing, as previously found in adults with an artificial script (Yoncheva et al., 2010, 2015). Words taught globally as whole visual shapes induced bilateral responses, while control letter strings (words/pseudowords), triggering GP mappings, led to typical LH responses. Importantly, this pattern was modulated by reading level and was present only in poor readers, which strongly suggests that the difficulty in automatizing GP mappings induced an increased reliance on an alternative visual strategy. Given that our data were collected in the context of a mixed approach and after only 2 months of school instruction, we conclude that the use of a global method very rapidly exposes children to a deviant reading strategy, which might impede the acquisition of reading especially in children having difficulties to acquire GP mappings. Therefore, in agreement with the view predominantly defended, our data is a strong argument in favor of the use of strict alphabetical approaches.

## Material and Methods

### Participants

First-grade children (N=44) from two different Belgian schools (20 boys, mean age = 6.08 years; range = 5.11-7.11, 41 right-handed) were tested at the very beginning of formal reading instruction (in the first trimester of grade 1, i.e., in November-December). Two of these children were excluded because of abnormal performances in behavioral tests (see below). All children had normal or corrected-to-normal vision. They were tested after the parents gave a written informed consent for a study approved by the Biomedical Ethical Committee of the University of Louvain. They were unaware of the goal of the study and that a change of stimulus type occurred at a periodic rate during stimulation. The testing took place in a quiet room of the school in two or more sessions (EEG, behavioral).

### Behavioral testing

General cognitive functions and reading ability were assessed by means of standardized tests and subtests: nonverbal intelligence (CPM; Raven, 1998), selective attention (TEA-Ch; Manly, Robertson, Anderson & Mimmo-Smith, 2004), vocabulary production (N-EEL; Chevrie-Muller & Plaza, 2001) and reading of single letters, syllables, regular words, irregular words, pseudo-words (BELO; George & Pech-Georgel, 2012), BALE; Jacquier-Roux, Lequette, Pouget, Valdois, & Zorman, 2010). In order to identify outliers within the distribution of the current sample, individual Z scores were computed for each general cognitive function. One child was excluded because of scores lower than 2 standard deviations in all general cognitive functions. Another child was excluded because of a medicated attentional disorder. Descriptive statistics of included children (table 1) highlight a great heterogeneity of reading scores within the group.

### EEG testing

#### Stimuli

Four categories of 20 stimuli were used for this experiment (Fig. 1.): pseudofonts (PF), words learned by a global method at school (GW) and control words (W) or pseudowords (PW). The natural school context of learning provided us with the “global words”. We chose 4- and 5-letters words (N=10 of each) from the « words’ box » that teachers had provided children at the beginning of the school year and which contained the words learnt item by item by mapping the whole-word form to its phonological counterpart. Control words were words for which children did not receive any explicit instruction and they were selected from the Manulex database (Lété, Sprenger-Charolles, & Colé, 2004) to match global words in lexical frequency, bigram frequency, orthographic neighborhood density and in number of letters (four or five). W and GW did not differ in *frequency estimated for grade 1* (t(38)=-0.89; *P* = 0.380); W = 79.40 ± 38.98 SD, GW = 102.00 ± 106,96 SD), *standard frequency index* (t(38)=0.60; *P* = 0.552); W = 65.35 ± 2.68 SD, GW = 64.56 ± 5.20 SD), *estimated frequency of use* (t(38)=-0.69; *P* = 0.495); W = 400 per million ± 197.68 SD, GW = 486 per million ± 525.18 SD), *bigram frequency* ((t(38)=-0.36; *P* = 0.971); W = 8390.10 ± 4261.80 SD, GW = 8440.95 ± 4601.25 SD) or *orthographic neighborhood density* ((t(38)=0.73; *P* = 0.467); W = 5.55 ± 4.46 SD, GW = 4.50 ± 4.57 SD). Pseudowords were pronounceable letter strings which respected the phonological rules in French. They were built one by one on the basis of the words: first and last bigrams (adjacent letters) of the 4-letters words were rearranged (e.g., the words “joli” (“cute”) and “fête” (“party”) could give rise to the pseudowords ‘’jote’’ and ‘’fêli’’) and the letters position of the 5-letters words was changed (e.g., the words “route” (“street”), “table” (“table”) and “forêt” (“forest”) give rise to the pseudowords “reuto”, “ablet” and “frêto”). Pseudowords were matched with all words (W and GW) in bigram frequency, identity of letters and in number of letters (four or five). Pseudowords did not differ in *bigram frequency* (8141.15 ± 3491.40 SD) from words (8390.10 ± 4261.80 SD; t(38)=0.20; *P* = 0.841) or global words (8440.95 ± 4601.25 SD; t(38)=0.23; *P* = 0.818). After the experiment, we assessed how many letters were used in the control letter strings that were also recognized by children during the behavioral letter recognition task. On average, children knew 14/18 (mean accuracy = 79.37% ± 19.80%, min = 27.78%, max = 100%) letters used in our stimuli (60% of the children knew 15/18 letters or more). Pseudofont stimuli were also built one by one on the basis of all the words (W and GW): each word was vertically flipped and its letters were segmented into simple features by using Adobe Photoshop. These segments were then rearranged to form pseudoletters, respecting the total number of characters (four or five) and the overall size (width χ height) of the original word (Lochy et al., 2015, 2016). Pseudoletters thus contained junctions, ascending/descending features and close-up shapes. Therefore, each word (W or GW) had a corresponding pseudoword and pseudofont, containing the exact same amount of black-on-white contrast, so that all conditions were comparable in terms of low-level visual properties.

These different stimuli allowed to create three conditions (Fig.1.). In each condition, base stimuli were pseudofonts (PF), and target stimuli were either global words (PF-GW condition), control words (PF-W condition) or pseudowords (PF-PW condition). Stimuli were presented centrally in Verdana font with a height between 47 and 77 pixels and a width between 103 and 271 pixels, depending on the shape of the individual letters. At a viewing distance of 1 m with a screen resolution of 800 χ 600 pixels and a refresh rate of 60 Hz, stimuli ranged from 2.69 to 7,07 (width) and 1.32 to 2.18 (height) degrees of visual angle.

#### Procedure

The stimulation procedure was very similar as in our previous FPVS-EEG studies on word recognition (Lochy et al., 2015, 2016). Each stimulation sequence started with a fixation square displayed for 2-5 s (randomly jittered between sequences), after which the stimulation gradually faded-in during 2 s (progressive increasing modulation depth from 0% maximum contrast level to 100%). Then, the sequence of stimulation was presented for 40 s, after which the stimulation faded-out during 2 s. This procedure was used to avoid abrupt eye-movements or blinks at the beginning or near to the end of a sequence. Stimuli were presented by means of sinusoidal contrast modulation at a base frequency rate of 6 Hz (i.e., one item every 166.66 ms, from a grey background to full contrast and back in 166.66 ms thus, each item reached full contrast at 83 ms). Given that the stimulus can be recognized at low contrast (i.e., 20% or less), the actual duration of stimulus visibility was close to 140 ms. Every fifth stimulus (1/5) of the sequence (frequency of 1.2 Hz thus, every 833 ms), a global word (PF-GW sequence) or a control letter strings (PF-W or PF-PW sequence) was presented. Stimuli were presented with a software (Sinstim) running over a JavaScript (Java SE Version 8). Each condition was repeated three times. Considering a total of 40 s (sequence duration) × 3 (repetitions) × 3 (conditions), 6 min of stimulation were presented in total. There was a pause of approximately 30 s between each sequence, which was initiated manually to ensure low-artifact EEG signals.

During the stimulation, children continuously fixated a central square and were instructed to press the space bar when they detected any brief (200 ms) color change of the fixation square (blue to yellow; six changes randomly timed per sequence) (see video 1). This orthogonal task was included to maintain a constant level of attention throughout the entire stimulation (see Lochy et al., 2015). Children performed this task almost at ceiling (95.93% ± 6.46 SD accuracy), showing high attention to the stimulation. There were no significant differences between conditions with respect to accuracy [F(2,78) < 1], or response time [F(2,78) < 1].

**Video 1.** 22 s excerpt of a stimulation sequence, showing pseudofonts at 6Hz, with (french) words (PF-W condition) appearing every five items (i.e., 1.2Hz).

#### Acquisition

During EEG recording, children were seated comfortably in a quiet room in the school at a distance of 1 m from the computer screen. EEG signal was acquired at 1,024Hz by using a 32-channel Biosemi Active II system (Biosemi, Amsterdam, Netherlands), with electrodes including standard 10–20 system locations. The magnitude of the offset of all electrodes, referenced to the common mode sense, was held below 50 mV.

#### Preprocessing

All EEG analyses were carried out by using Letswave 5.c (http://nocions.webnode.com/letswave) and Matlab 2014 (The Mathworks) and followed procedures validated in several studies using letter strings or faces and objects stimuli (see e.g., Retter & Rossion, 2016). After band-pass filtering between 0.1 and 30 Hz, EEG data were segmented to include 2 s before and after each sequence, resulting in 44 s segments. Data files were then downsampled to 256 Hz to reduce file size and data processing time. Artifact or noisy channels were replaced by using linear interpolation. All channels were referenced to the common average. EEG recordings were then segmented again from stimulation onset until 39.996 s, corresponding exactly to 48 complete 1.2Hz cycles within stimulation. This is the largest amount of complete cycles of 833 ms at the target frequency (1.2 Hz) within the 40 s of stimulation period.

#### Frequency domain analysis

To reduce EEG activity that is not phase-locked to the stimulus, the three repetitions of each condition were averaged in the time domain for each individual participant. Then, to convert data from the time domain into the frequency domain, a Fast Fourier Transform (FFT) was applied to these averaged time windows and normalized amplitude spectra were extracted for all channels. Thanks to the long time-window (39.996 s), this procedure yields EEG spectra with a high frequency resolution (1/39.996 s = 0.025Hz), increasing SNR (Regan, 1989; Rossion, 2014) and allowing unambiguous identification of the response at the exact frequencies of interest (i.e., 6Hz for the base stimulation rate and 1.2Hz and its harmonics for the target stimulation rate). All of the responses of interest, and thus all the potential differences between conditions, can be concentrated in a discrete frequency band around the stimulation frequency. This frequency band occupies a very small fraction of the total EEG bandwidth. In contrast, biological noise is distributed throughout the EEG spectrum, resulting in a SNR in the bandwidth of interest that can be very high (Regan, 1989; Rossion, 2014). To estimate SNR across the EEG spectrum, amplitude at each frequency of interest (bin) was divided by the average amplitude of 20 surrounding bins (10 on each side) (Liu-Shuang, Norcia, & Rossion, 2014).

To quantify the responses of interest in microvolts, the average voltage amplitude of the 20 surrounding bins (i.e., the noise) was subtracted out (e.g., Dzhelyova & Rossion, 2014; Retter & Rossion, 2016) (baseline-subtracted amplitudes). To assess the significance of the responses at the target frequency and harmonics, and at the base rate and harmonics, Z scores were computed at every channel on the grand averaged amplitude spectrum for each condition (e.g., Liu-Shuang et al., 2014; Lochy et al., 2015). Z scores larger than 2.58 (P < 0.01, one-tailed, signal > noise) were considered significant. A conservative threshold was used because the response was evaluated on all channels (although we expected responses at posterior channels), and distributed on several harmonics as in previous studies using the same approach with letters strings (Lochy et al., 2015, 2016) or faces (Liu-Shuang et al., 2014). An identical number of harmonics was selected across all conditions and electrodes based on the condition in which the highest number of consecutive harmonics was significant on any electrode (in total, 4 harmonics for target responses and 5 harmonics for base responses). Finally, to quantify the periodic response distributed on several harmonics, the baseline subtracted amplitudes of significant harmonics (except the base stimulation frequency) were summed for each participant (see Retter & Rossion, 2016 for validation of the procedure).

## Acknowledgements

We thank the schools and children for participation.

